# Haplocheck: Phylogeny-based Contamination Detection in Mitochondrial and Whole-Genome Sequencing Studies

**DOI:** 10.1101/2020.05.06.080952

**Authors:** Hansi Weissensteiner, Lukas Forer, Liane Fendt, Azin Kheirkhah, Antonio Salas, Florian Kronenberg, Sebastian Schoenherr

## Abstract

Within-species contamination is a major issue in sequencing studies, especially for mitochondrial studies. Contamination can be detected by analysing the nuclear genome or by inspecting the heteroplasmic sites in the mitochondrial genome. Existing methods using the nuclear genome are computationally expensive, and no suitable tool for detecting contamination in large-scale mitochondrial datasets is available. Here we present haplocheck, a tool that requires only the mitochondrial genome to detect contamination in both mitochondrial and whole-genome sequencing studies. Haplocheck is able to distinguish between contaminated and real heteroplasmic sites using the mitochondrial phylogeny. By applying haplocheck to the 1000 Genomes Project data, we show (1) high concordance in contamination estimates between mitochondrial and nuclear DNA and (2) quantify the impact of mitochondrial copy numbers on the mitochondrial based contamination results. Haplocheck complements leading nuclear DNA based contamination tools, and can therefore be used as a proxy tool in nuclear genome studies.

Haplocheck is available both as a command-line tool at https://github.com/genepi/haplocheck and as a cloud web-service producing interactive reports that facilitates the navigation through the phylogeny of contaminated samples.

## Introduction

The human mitochondrial DNA (mtDNA) is an extranuclear DNA of ∼16.6 kb length (Andrews et al. 1999). It is inherited exclusively through the maternal line facilitating the reconstruction of the human maternal phylogeny and female (pre-)historical demographic patterns worldwide. The strict maternal inheritance of mtDNA results in a natural grouping of haplotypes into monophyletic clusters, referred to as haplogroups (Kivisild et al. 2006; Kloss-Brandstätter et al. 2011). Furthermore, second generation sequencing enables the detection of heteroplasmy over the complete mitochondrial genome. Heteroplasmy is the occurrence of at least two different haplotypes of mtDNA in the investigated biological samples (e.g. cells or tissues). Depending on the sequencing coverage, heteroplasmic positions are reliably detectable down to the 1% variant level (Weissensteiner et al. 2016; Ye et al. 2014). In recent years, the issue on apparent heteroplasmy in mitochondrial data and data interpretation was addressed by several studies (Bandelt and Salas 2012; He et al. 2010; Ye et al. 2014; Just et al. 2014) resulting in a comprehensive review on the quality of mtDNA data derived from sequencing studies (Just et al. 2015). It has been shown that certain studies can overestimate the presence of heteroplasmy, which can often be explained by external or cross-contamination (Yao et al. 2007; Just et al. 2014, 2015; Brandhagen et al. 2020; Yin et al. 2019), artificial recombination (Bandelt et al. 2004), artifacts, index hopping (Van Der Valk et al. 2019) or analysis software inconsistencies. Sample contamination is still a major issue in both nuclear DNA (nDNA) and mtDNA sequencing studies that must be prevented to avoid mistakes as it occurred with Sanger sequencing studies in the past (Salas et al. 2005). Due to the accuracy and sensitivity of second generation sequencing combined with the availability of improved computational models, within-species contamination is traceable down to the 1% level in whole-genome sequencing (WGS) studies (Jun et al. 2012).

Several approaches exist to detect contamination in mtDNA sequencing studies. We previously showed that a contamination approach based on the co-existence of phylogenetically incompatible mitochondrial haplotypes observable as heteroplasmy is feasible (Weissensteiner et al. 2016), which has also been demonstrated by others (Avital et al. 2012; Li et al. 2010, 2015). Other methods, such as a Galaxy-based approach (Dickins et al. 2014) facilitates the check for contamination by building neighbor joining trees. Mixemt (Vohr et al. 2017) incorporates the mitochondrial phylogeny and estimates the most probable haplogroup for each sequence read; the computational expensive algorithm implemented in Mixemt reveals advantages for contamination detection of several haplotypes within one sample and is independent of variant frequencies. For ancient DNA studies, schmutzi (Renaud et al. 2015) uses sequence deamination patterns and fragment length distributions to estimate contamination. Additionally, specific lab-protocols were designed for eliminating contamination, including double-barcode sequencing approaches (Yin et al. 2019).

For contamination detection in mitochondrial studies, cross-contamination using the nuclear genome is often investigated (Wei et al. 2019; Ding et al. 2015; Yuan et al. 2020) by applying widely accepted software tools like VerifyBamID (Zhang et al. 2020; Jun et al. 2012). Nevertheless, it becomes apparent that a tool for mitochondrial studies is missing that is able to detect contamination using the mitochondrial genome. Furthermore, since mtDNA is also present hundred to several thousand-fold per cell depending on cell-type, also WGS datasets specifically targeting the autosomal genome result in a high coverage over the mitochondrial genome. We hypothesize that the nDNA contamination level might be estimated by looking only at the mitochondrial genome.

In this paper, we systematically evaluate the approach of using the mtDNA phylogeny for contamination detection and present haplocheck, a tool to detect contamination in mtDNA and WGS studies. In general, haplocheck works by identifying heteroplasmic sites (both real and artificial due to contamination) starting both from CRAM, BAM or VCF files. Using the mitochondrial phylogeny and the concept of haplogroups, haplocheck is able to report contamination estimates by identifying two stable haplogroups within one sample. Overall, this work should demonstrate the merits of the mitochondrial genome as an instrument for rapid contamination detection in sequencing studies and presents a tool that takes advantage of a solid well-known mitochondrial phylogeny.

## Methods

Haplocheck takes either CRAM, BAM or VCF files as an input. For files in the CRAM or BAM format, an initial step to detect homoplasmic and heteroplasmic sites using a Maximum Likelihood (ML) function (Ye et al. 2014) is executed and final sites are reported in the variant call format (VCF). Heteroplasmic sites are then split by their allele frequency (AF) into a major and minor profile. A profile consists of all detected homoplasmic and the corresponding fraction of each heteroplasmic variant. Heteroplasmic fractions with an AF >=50% are added to the major profile, otherwise to the minor profile. The haplogroup for each profile is then determined using Haplogrep2 (Weissensteiner et al. 2016b). Using the mitochondrial phylogeny, the phylogenetic distance (i.e. number of nodes between the two haplogroups) is calculated. The identification of two stable haplogroups allows haplocheck to report the contamination level for each sample.

Three different scenarios need to be considered for contamination detection based on the mitochondrial phylogeny. First, two haplotypes branch into two different nodes: a major haplotype with heteroplasmy level x and a minor haplotype with heteroplasmy level 1-x (Figure 1A), whereas H1a1 represents the Last Common Ancestor (LCA) for both haplotypes. Second, if heteroplasmic sites are only identified in the major haplotype, the minor haplotype H1a1 is defined as the LCA (Figure 1B). Third, if heteroplasmic sites are only present in the minor haplotype, the major haplotype H1a1 defines the LCA (Figure 1C).

**Figure 1:**
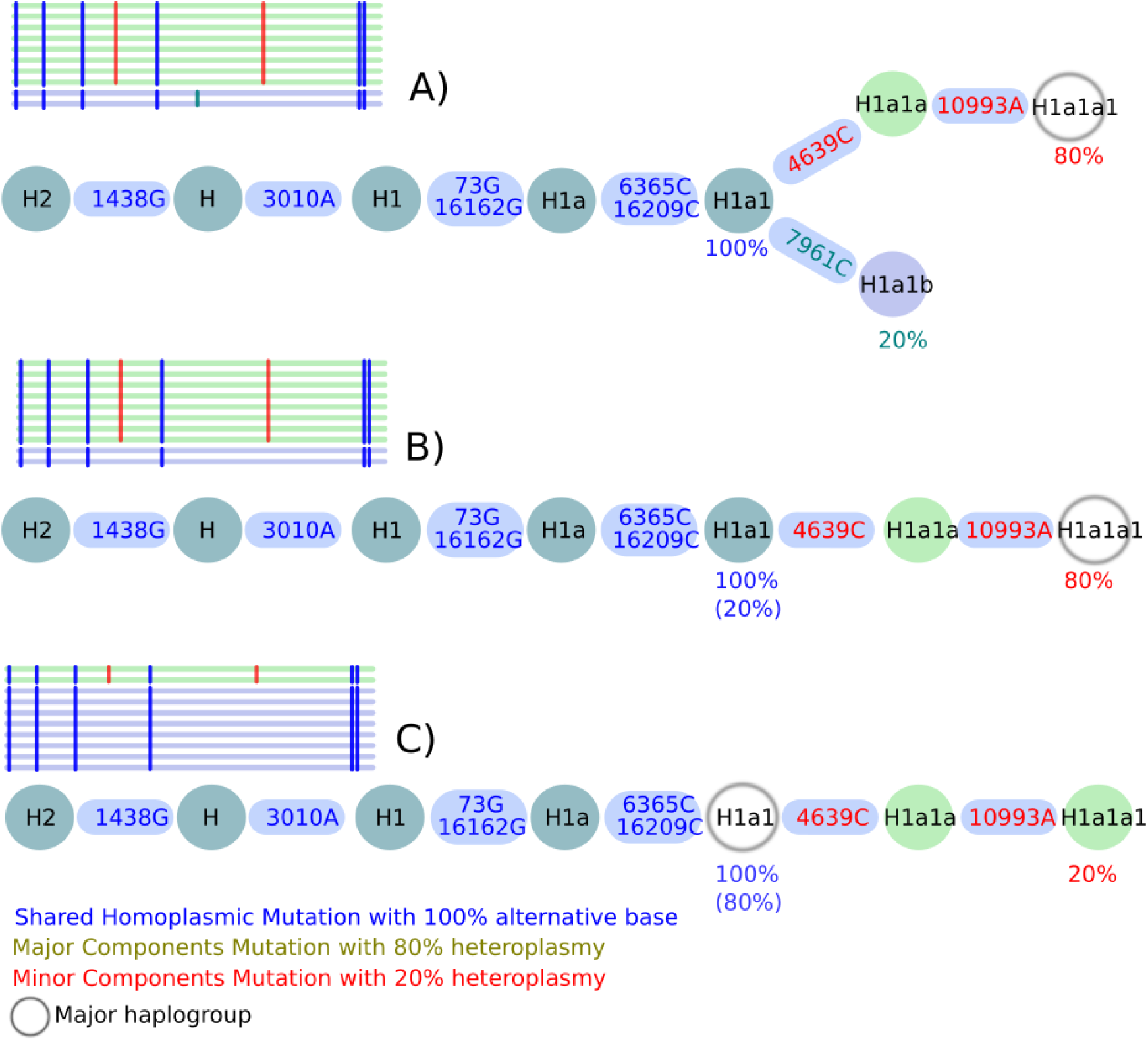
All possible contamination scenarios. Here, a contamination level of 20% is shown in all three scenarios A) to C). Shared polymorphisms of two haplotypes are included in a single branch, whereas the split into two branches displays the different lineage haplotypes. **A)** Shared mutations defining H1a1 (Last Common Ancestor, LCA) are present at 100%, while 7961C is present only at 20% defining the minor haplogroup H1a1b, whereas 4639C and 10993A is present at 80% defining the major haplogroup H1a1a1. **B)** A mixture of two haplotypes within a single lineage but of different lineage depths (minor haplotype H1a1 and major haplotype H1a1a1) is observed if no minor haplotype can be found. **C)** A mixture of two haplotypes within a single lineage but of different lineage depths (minor H1a1a1 and major H1a1) is observed if the minor haplotype results in a haplogroup. Shared homoplasmic sites facilitate the identification of the branching pattern in all three scenarios and improving the overall haplogroup quality. The used notation for variants (e.g. 1438G) includes the mtDNA position (1438) followed by the actual base change (G).

### Heteroplasmic and Homoplasmic Variant Calling

For BAM or CRAM files, the overall performance of haplocheck relies on an accurate homoplasmic and heteroplasmic variant calling. Therefore, we previously developed mtDNA-Server (Weissensteiner et al. 2016) that allows the detection of heteroplasmic positions accurately down to 1%. The identification of heteroplasmic positions is based on a previously published method (Ye et al. 2014) that also includes several criteria for calling heteroplasmy such as (a) base quality >= 20, (b) >10 × depth per strand, (c) 1% minor allele frequency on each strand and (d) a log-likelihood ratio (LLR) of >=5. LLR represents the ratio between the estimated frequency of the major allele within the ML function of the heteroplasmy and the homoplasmy model.

For this work, we re-implemented mtDNA-Server as a standalone module (https://github.com/seppinho/mutserve) and integrated it into haplocheck. Detected heteroplasmic positions are reported in VCF format as heterozygous genotypes (GT) using the AF tag for the estimated contamination level. Although the term genotype makes sense in autosomal diploid scenarios, we use it here to refer to mtDNA variation patterns that resemble a genotype status.

For homoplasmic positions, the final genotype GT ({A,C,G,T}) is detected using all input reads (reads) and calculating the genotype probability P using Bayes’ Theorem P(GT|reads) = P(reads|GT) × P(GT) / P(reads). To calculate the prior probability P(GT), we used the 1000 Genomes Phase 3 VCF file and calculated the frequencies for all sites using vcftools (Danecek et al. 2011). To compute P(reads|GT), we calculate the sequence error rate (e_i_ = 10^−Qi/10^) for each base *i* of a read, whereas *Q* is the reported quality value. For each genotype GT (GT ∈ {A,C,G,T}) of a read, we determine the genotype likelihood by multiplying 1-e_i_ in case the base of the read r_i_ = GT and e_i_/3 otherwise over all reads (Ding et al. 2015). The denominator P(reads) is the sum of all four P(reads|GT).

### Contamination Detection Model

The contamination model within haplocheck requires a VCF file as an input and includes steps for (a) splitting homoplasmic and heteroplasmic sites into two haplotype profiles, (b) haplogroup classification for each haplotype profile, and (c) applying quality-control criteria. Homozygous genotypes for the alternate alleles (ALT; i.e. homoplasmic sites) are added to both haplotypes and heterozygous genotypes (i.e. real and artificial heteroplasmic sites) and are split using the AF tag. Since mutserve always reports the AF of the non-reference allele, the split method applies the following rule: In case a GT 0/1 (e.g. Ref: G, ALT: C) with an AF of 0.20 is included, the split method defines C as the minor allele, 0.2 as the minor level and 0.8 as the major level. In case a GT 0/1 (e.g. Ref: G, ALT: C) with an AF of 0.80 is included, the C defines the major allele. If no reference allele is included (e.g. 1/2), we use the first allele as the major allele and assign the included AF to that allele.

For haplogroup classification, we use HaploGrep2 based on Phylotree 17 (van Oven and Kayser 2009), which has been refactored as a module and integrated directly into haplocheck. As a result, Haplogrep2 reports the haplogroup of both the major and minor haplotype. For each analyzed sample, the LCA is calculated, which is required to estimate the final contamination level and to calculate the distance between the two haplotypes. Therefore, we traverse Phylotree from the rCRS reference to each node. The LCA is determined by starting at the final node of haplotype 1 (h1) and by iterating back until the reference (rCRS) is reached. Then, we iterate back to rCRS for haplotype 2 (h2) until the first node included in h1 is identified. This node then defines the LCA of both haplotypes. Only heteroplasmic positions starting from the LCA and showing a phylogenetic weight >5 are included. The phylogenetic weight describes the frequency of each mutation in Phylotree and is scaled from 1 to 10 in a non-linear way. Variants with a high occurrence in Phylotree are assigned a small phylogenetic weight. Furthermore, back mutations (i.e. mutation changes back to the rCRS reference within a specific haplogroup) and deletions on heteroplasmic sites are ignored by haplocheck.

Using all previous information, we finally estimate the contamination level for samples fulfilling the following three quality control criteria: (a) >=2 heteroplasmic variants starting from the LCA, (b) >=0.5 haplogroup quality for each haplotype (calculated by HaploGrep2 using the Kulczynski metric) and (c) phylogenetic distance betcomponentween both haplotypes of >=2. The median level of all detected heteroplasmic sites reaching the described criteria is calculated for both haplotypes (h1 and h2) independently. Haplocheck reports the median level of the minor haplotype as the final contamination level.

### Report

Haplocheck produces a tab-delimited text file and an interactive HTML report. For each sample, haplocheck determines the final contamination status, the contamination level and several quality metrics such as the phylogenetic distance. Additionally, a graphical phylogenetic tree is generated dynamically for each sample including the path from the rCRS to the two final haplotypes. This allows the user to manually inspect edge cases, visualize the contamination graphically or analyze the source of contamination (see Supplemental Figure S1).

## Results

Haplocheck can be used as a standalone command-line tool or as a cloud web service. For both scenarios, the same workflow is applied. The Cloudgene framework (Schönherr et al. 2012) is utilized to provide the workflow as-a-service to users, which is also used for large-scale genetic services like the Michigan Imputation Server (Das et al. 2016) and the mtDNA-Server (Weissensteiner et al. 2016) that greatly improves user experience and productivity.

### Evaluation

To test the performance of haplocheck within mtDNA and WGS studies, we simulated several data sets. In a first step, we created wet-lab mixtures of two mitochondrial samples to validate the variant calling with mutserve. The mixtures were as follows: M1 - 1:2 (50%), M2 - 1:10 (10%), M3 - 1:50 (2%), M4 - 1:100 (1%), M5 - 1:200 (0.5%, created in-silico). All mixtures and the two initial samples have been then sequenced on an Illumina HiSeq system. We analyzed the original samples (coverage 60,000 ×) and downsampled them accordingly. Table 1 summarizes our findings and shows that a coverage between 600 - 900 is required, to detect contamination down to 1%. For example, the 1000 Genomes Project low-coverage sequence data already include an average coverage of 1797 of the mitochondrial genome (1000 Genomes Project Consortium 2015).

**Table 1:**
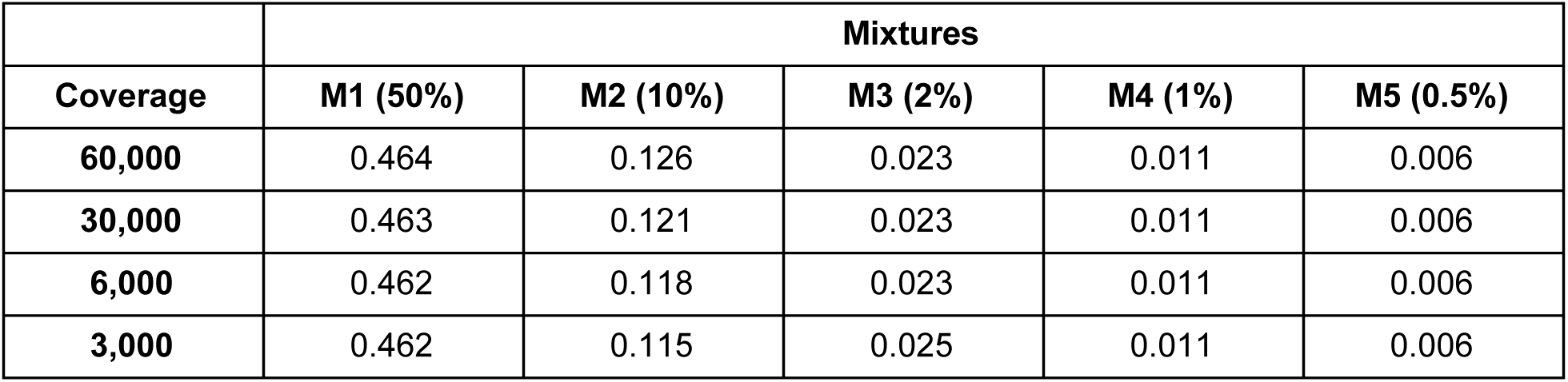
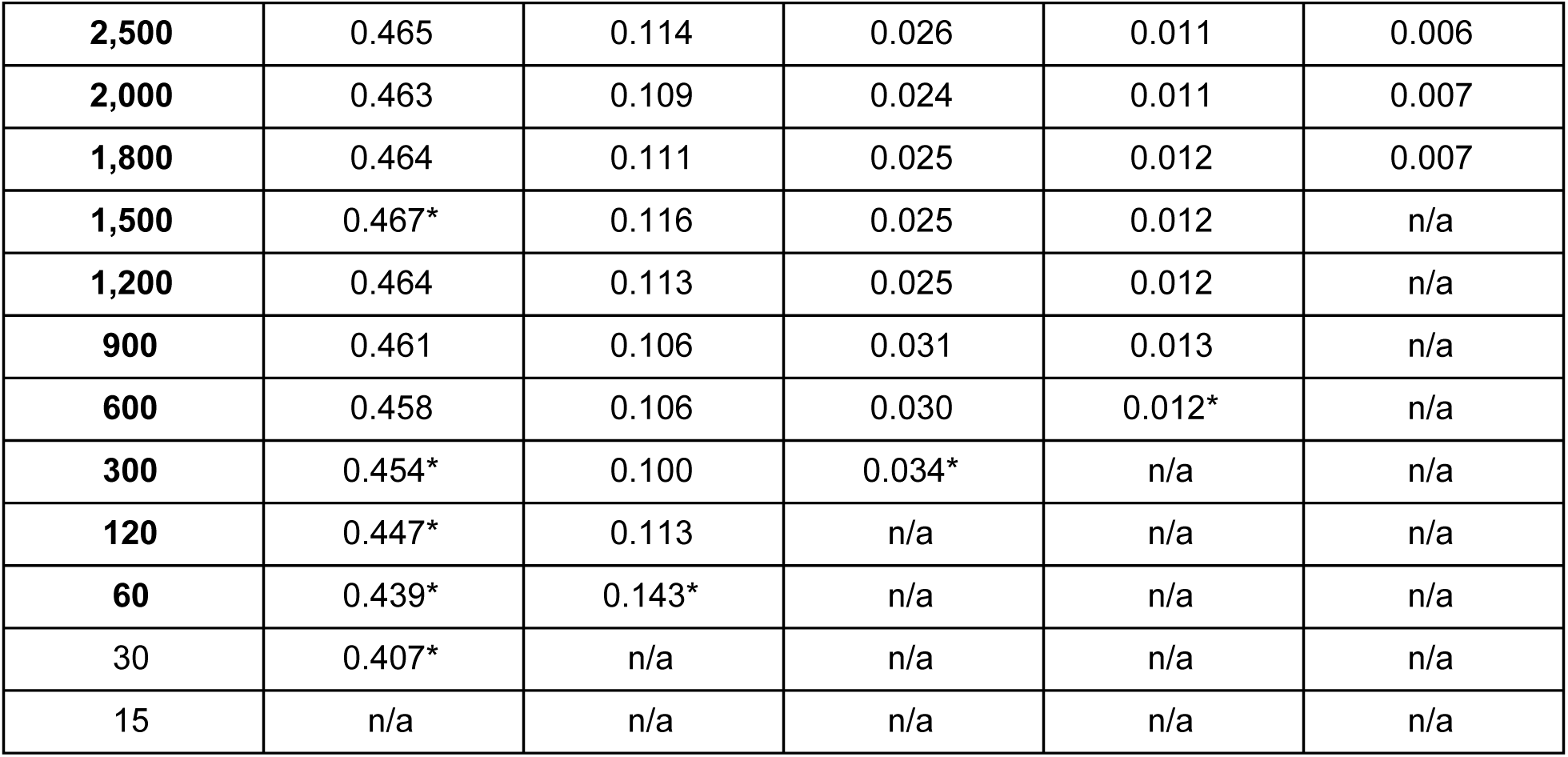
Four wet-lab mixtures (M1-M4) and 1 in-silico mixture (M5) have been analyzed using haplocheck with varying coverage. The columns “M1 - M5” indicate the mixture levels, “Coverage” indicates the downsampled coverage. Each cell in the table includes either the actual detected contamination level reported by haplocheck or n/a in case the contamination could not be detected by haplocheck. The asterisk (*) indicates that the detected haplotypes differed from the expected haplotypes, since not all variants were detected in the mixture.

|  | Mixtures |  |  |  |  |
| --- | --- | --- | --- | --- | --- |
| Coverage | M1 (50%) | M2 (10%) | M3 (2%) | M4 (1%) | M5 (0.5%) |
| 60,000 | 0.464 | 0.126 | 0.023 | 0.011 | 0.006 |
| 30,000 | 0.463 | 0.121 | 0.023 | 0.011 | 0.006 |
| 6,000 | 0.462 | 0.118 | 0.023 | 0.011 | 0.006 |
| 3,000 | 0.462 | 0.115 | 0.025 | 0.011 | 0.006 |
| **2,500** | 0.465 | 0.114 | 0.026 | 0.011 | 0.006 |
| **2,000** | 0.463 | 0.109 | 0.024 | 0.011 | 0.007 |
| **1,800** | 0.464 | 0.111 | 0.025 | 0.012 | 0.007 |
| **1,500** | 0.467* | 0.116 | 0.025 | 0.012 | n/a |
| **1,200** | 0.464 | 0.113 | 0.025 | 0.012 | n/a |
| **900** | 0.461 | 0.106 | 0.031 | 0.013 | n/a |
| **600** | 0.458 | 0.106 | 0.030 | 0.012* | n/a |
| **300** | 0.454* | 0.100 | 0.034* | n/a | n/a |
| **120** | 0.447* | 0.113 | n/a | n/a | n/a |
| **60** | 0.439* | 0.143* | n/a | n/a | n/a |
| 30 | 0.407* | n/a | n/a | n/a | n/a |
| 15 | n/a | n/a | n/a | n/a | n/a |

We further generated sequencing data mixtures by using the ART-NGS read simulator (Huang et al. 2012). The four generated mixtures differ in the contamination level (0.5% - 50 %), coverage (between 10 × - 5000 ×) and phylogenetic distance between the two mixed samples (3 - 23 nodes). The results were highly concordant with the wet-lab mixtures presented in Table 1 and show that haplocheck is able to detect contamination accurately even for samples including haplotypes with a short phylogenetic distance (see Supplemental Table S1).

In a second step, we evaluated the performance of haplocheck as a proxy for estimating the nDNA contamination level. Therefore, we generated four whole-genome in-silico samples from two random 1000 Genomes samples showing no signs of contamination based on the VerifyBamID score. For each sample, four different mixtures between 0.5% - 10% have been created and analyzed using both VerifyBamID2 (nDNA) and haplocheck (mtDNA). To analyze the impact of the mitochondrial copy number (mtCN), samples with different amounts of mtCN were chosen from the 1000 Genomes Project. Table 2 summarizes the findings, whereby each cell includes the average delta between the calculated and the expected value for all four different mixtures per sample (see also Supplemental Table S2). Levels obtained from VerifyBamID2 and haplocheck correlate if the copy number (CN) for each haplotype in the mixture is similar (see Mixture 1 and 2). Values obtained from Mixture 3 still correlate, since the main haplotype shows a higher mtCN and is therefore unaffected by the lower mtCN of haplotype 2. In a worst-case scenario (Mixture 4), where the main haplotype has a lower mtCN and the minor haplotype a higher mtCN, the values between haplocheck and VerifyBamID2 differ substantially.

**Table 2:** Four different mixtures have been created and the average delta between expected and calculated contamination level reported. Each average delta consists of four different mixtures (1-10%) and has been calculated for VerifyBamID2 using a different set of markers as well as haplocheck. Haplocheck works well as a proxy for the first two sample mixtures, but differs as expected in substantially uneven mtCN between the main haplotype (low mtCN) and the second haplotype (high mtCN).

|  | mtCN Ratio | VerifyBamID2 |  |  |  | Haplocheck |
| --- | --- | --- | --- | --- | --- | --- |
|  |  | HGPD_100K | HGPD_10K | 1000G_100K | 1000G_10K | Phylotree 17 |
| **Mixture 1** | 1:1 | -0.85% | -0.51% | -0.34% | 0.11% | 0.45% |
| **Mixture 2** | 1:0.8 | -0.26% | -0.08% | -0.49% | -0.12% | 1.32% |
| **Mixture 3** | 10:1 | -0.66% | -0.66% | -0.50% | -0.61% | -3.70% |
| **Mixture 4** | 1:10 | -0.03% | -0.06% | -0.22% | -0.36% | 20.85% |

Nevertheless, a drastic shift in the copy number as in mixture 4 is atypical for a sequencing project. In (Zhang et al. 2017) the copy number of 1,500 women with age 17-85 have been analyzed and show that most samples ranging from 100 - 300 (mean 169, DNA source whole-blood). In (Fazzini et al. 2019) the mtCN has been analysed in a cohort of 4,812 chronic kidney disease patients showing also only moderate differences (mean 107.2, sd 36.4, DNA source whole-blood).

In a third step, we created and analyzed in-silico data by mixing random genotype profiles from the currently best available mtDNA phylogeny derived from Phylotree Build 17. The overall performance of haplocheck heavily depends on a good classification of samples into haplogroups even from noisy variant calling data sets. We initially created input profiles for each displayed haplogroup, amounting to 5,426 profiles in total. Each input profile consists of a list of polymorphisms from the tree reference (rCRS) to the actual node (or haplogroup). Our test data consists of 500,000 unique mixtures of pairwise haplogroup profiles derived from the overall phylogeny comprising of 5,500 haplogroups (250,000 contaminated, 250,000 not-contaminated samples) and 100,000 mixtures from the haplogroup H-subtree, including 977 haplogroups. The generation of in-silico data from the H-subtree allows us to test the performance of samples showing a smaller phylogenetic distance.

To account for noisy input data, we artificially added random variants to each input profile. This has been done by removing expected variants from the input profile and adding random variants available within Phylotree. The amount of noise varies from 0 - 8 variants for each mixture. The proportion of added *versus* removed variants is calculated randomly. To make it further restrictive, we only added phylogenetic relevant variants from Phylotree. Variants that are not present in Phylotree (i.e. so far unknown in the phylogeny) would not affect the contamination estimation. Finally, 3 datasets (noise 0, 4, 8) derived from 2 different trees (complete tree, haplogroup H subtree) have been generated, each consisting of 500,000 and 100,000 mixtures respectively. F1-Score (defined as (2 × precision × sensitivity) / (precision + sensitivity)) has been calculated for each mixture to analyze the overall accuracy of haplocheck.

To determine the best haplocheck configuration regarding accuracy, we tested different setups for all 6 datasets. Each setup includes a different threshold for (1) the amount of major and minor heteroplasmic sites, (2) the minimum allowed phylogenetic distance between two profiles and (3) the haplogroup classification model (Kulczynski, Hamming, Jaccard). Figure 2 summarizes the 6 best setups that have been tested to determine the optimal trade-off between noise, haplogroup distance and the overall F1-Score. In our experiments, Setup 3 shows the best trade-off between haplogroup distance and overall accuracy. This setup allows us to detect contamination of samples with a phylogenetic distance of at least 2 and has been used as the final setup for the contamination method.

**Figure 2:**
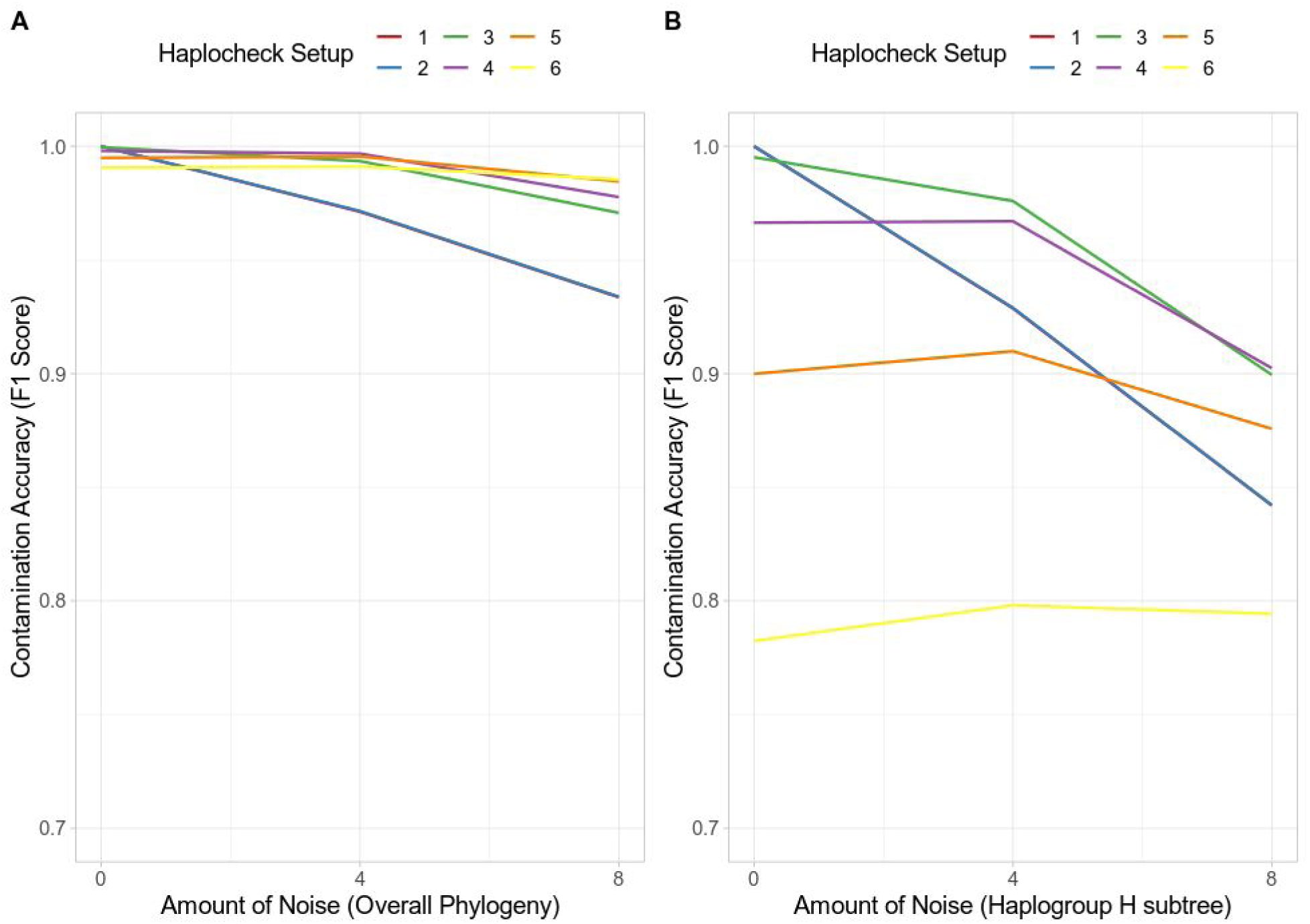
Tested haplocheck setups (=lines) to determine the best trade-off between noise and overall accuracy. Setup 3 (phylogenetic distance >= 2, amount of heteroplasmic sites >= 2, haplogroup quality > 0.5, Kulczynski Metric) shows the best trade-off for all 6 datasets. Each dataset consists of 500.000 mixtures (Overall Phylogeny) and 100.000 mixtures (Haplogroup H subtree) respectively. The x-axis includes the amount of noise, the y-axis the calculated F1-Score (scale from 0 to 1, where 1 equates to a perfect precision and recall).

Table 3 summarizes the F1 Score statistics for Setup 3. The result demonstrates that haplocheck is able to accurately detect contamination of two samples also in the case where noise is included in the input profiles and the distance between the two haplogroups is small.

**Table 3:** F1 Score for different noise categories using the finally chosen Setup 3. Noise 0 - Noise 8 includes the amount of added / removed variants from the input profile. The two experiments based on different trees (mixtures derived from the complete phylogenetic tree and mixtures derived from the haplogroup H subtree only) show that haplocheck is capable of detecting contamination accurately.

| **In-Silico Simulation**<br>Setup 3: Distance: 2; Heteroplasmies: 2, Kulczynski Metric |  |  |  |
| --- | --- | --- | --- |
| **Metric** | **Noise 0** | **Noise 4** | **Noise 8** |
| F1 Score Complete Phylogenetic Tree | 0.999 | 0.993 | 0.971 |
| F1 Score H Phylogenetic Tree | 0.995 | 0.976 | 0.899 |

### Contamination Detection in the 1000 Genomes Project

To evaluate haplocheck on a WGS study, we extracted the mtDNA genome reads (labeled as chromosome MT) from the 1000 Genomes Project^1^ (Phase 3), resulting in a sample size of 2,504 and a total file size of 95 GB. As an initial check, we compared variants detected by mutserve to the official 1000 Genomes data release using callMom (https://github.com/juansearch/callMom) and determined the haplogroup using HaploGrep2. Overall, 98 % of the samples (n = 2,504) result in an identical haplogroup (See Supplemental Figure S2). The downloaded BAM files have then been used as an input for haplocheck to test for contamination. Based on the mitochondrial genome, 5.07% (127 of 2,504) of all samples show signs of contamination on mtDNA (see Supplemental Table S3). Since the performance of haplocheck as a proxy for nDNA is dependent on the mtCN, we also looked at the tissue source used for DNA extraction. As depicted in Table 4 and Supplemental Figure 3, there is a significant difference in the mtCN of the two tissue types used within the 1000 Genomes Project (p<2.2e-16, independent *t*-test). The mtCN has been inferred using the formula mtDNA coverage / nDNA coverage × 2 (Ding et al. 2015).

**Table 4.** Tissue Cell Types of all 2,504 samples from the 1000 Genomes Project. Significant differences in the mitochondrial copy number (mtCN) between 1000G samples can be seen. Each cell includes the absolute and relative number of samples. LCL: lymphoblastoid cell lines.

|  | Tissue Cell Type |  |  |
| --- | --- | --- | --- |
| 1000 Genomes Phase 3 (n= 2,504) | Blood | LCL | Not specified |
| Samples Absolute / Relative | 364 / 14.5% | 506 / 20.2% | 1634 / 65.3% |
| Mitochondrial Copy Number (mtCN) Mean | 49.3 | 747.1 | 566.9 |

Due to the different mtCN, we split the 1000 Genomes samples into two groups based on the mtCN and calculated the Pearson correlation coefficient (R) separately. Group 1 (mtCN >=300, n=2,004) shows a correlation of R=0.72 between the contamination levels of VerifyBamID2 and haplocheck, whereas the contamination levels reported by haplocheck are ranging from 0.8% to 4.8% (see Supplemental Table 4). Group 2 (mtCN <300, n=500) shows a correlation of R=0.31 and contamination levels reported by haplocheck are between 1.8% - 25.5% (see Supplemental Table 5).

As expected, samples with a higher mtCN (group 1) are less vulnerable to level differences between nDNA and mtDNA. Therefore, mtDNA contamination levels are in a very similar range compared to those observed by VerifyBamID2 (0-3%). Samples with a lower mtCN (group 2) are more vulnerable to mtCN differences. This is due to the fact that a contamination with a sample showing a higher amount of mtCN can affect the contamination level substantially. Therefore, group 2 shows a much higher discrepancy in the contamination level compared to VerifyBamID2.

As mentioned earlier, such a drastic shift in the copy number is atypical for a sequencing project. For studies showing only moderate differences in the mtCN, haplocheck can be used as an efficient nDNA proxy. For studies showing a wider range of mtCN due to different tissues, the mtDNA level can differ from the reported nDNA level.

In the last step, we looked at samples that have been excluded from the 1000 Genomes Project (nDNA contamination level >3% using VerifyBamID). In total, 4 samples have been excluded by VerifyBamID due to a high free mix (sequence-only estimate of contamination) and 7 samples due to a high chip mix (for estimating contamination or swap using sequence and array method). Haplocheck was able to identify these samples as contaminated with a correlation of 0.89 between nDNA and mtDNA (Supplemental Table S6).

### Nuclear DNA of mitochondrial origin

Nuclear DNA of mitochondrial origin (NUMTS) can either result in a coverage drop on mtDNA sites due to the alignment of mitochondrial reads to NUMTS or false positive heteroplasmy calls due to the alignment of NUMT reads to the mitochondrial genome (Maude et al. 2019). Approaches exist (Goto et al. 2011; Samuels et al. 2013) that exclude reads mapping to the nDNA but overall reduce coverage and may result in false negatives (Albayrak et al. 2016). In (Weissensteiner et al. 2016), we annotated mitochondrial sites coming from an NUMTS reference database (Li et al. 2012; Dayama et al. 2014), although limited to known NUMTS. For contamination detection with haplocheck, false positive heteroplasmic sites due to NUMTS are expected to only have a minor effect since they typically do not resemble the complete mitochondrial haplotypes. Nevertheless, sufficient coverage for the haplogroup defining variants is still required when dealing with NUMTS. In a study conducted by (Maude et al. 2019), an in-silico model has been set up to analyze the homology between mitochondrial variants and NUMTS. They show that 29 variants representing haplogroups A, H, L2, M, and U did not cause loss of coverage, nevertheless substantial loss of coverage has been identified for specific sites (e.g.G1888A, A4769G). In a recent work the presence of a mega-NUMT that could mimic contamination on mitochondrial haplogroup level is described (Balciuniene and Balciunas 2019). This indicates that in very rare cases, NUMTs could indeed resemble complete mitochondrial haplotypes and yield to a false positive contamination result (Salas et al. 2020; Wei et al. 2020). While we did not observe NUMT-related issues in the validation of the 1000 Genome Project, we can not entirely rule out eventual NUMTs effects on contamination detection.

### Runtime and Performance

Table 5 shows that our pipeline starting from BAM data scales linearly with the data size (i.e. sequence reads). For the complete 1000 Genomes Project data, the contamination estimate has been calculated within 8 hours and 58 minutes starting from BAM using a single core (Intel Xeon CPU 2.30GHz) and 2 GB of RAM.

**Table 5.** Haplocheck v1.1.3 runtime for different BAM files. Runtime includes both variant calling (using mutserve) and contamination detection. All tests have been executed using a single core (Intel Xeon Processor E5-2650 v3) and 2 GB of RAM.

| Runtime Haplocheck | File Size |
| --- | --- |
| 1 min 49 sec | 0.19 GB |
| 2 min 58 sec | 0.37 GB |
| 15 min 55 sec | 1.85 GB |
| 31 min 26 sec | 3.7 GB |
| 8 h 58 min | 95 GB (2,504 samples) |

Table 6 includes a runtime comparison for 26 samples of VerifyBamID2 (input WGS data, varying amounts of markers and cores) with haplocheck (input mtDNA) starting from BAM data.

**Table 6.** Haplocheck v1.1.3 runtime for 26 samples of the 1000 Genomes Project data. For haplocheck, runtime includes variant calling with mutserve and contamination detection. For VerifyBamID2, all autosomes have been analyzed with different sets of markers (10k and 100k), therefore resulting in a much larger data size. All tests have been executed on an Intel Xeon Processor E5-2650 v3 CPU using OpenJDK 8 for haplocheck.

|  | Haplocheck | VerifyBamID2 |  |
| --- | --- | --- | --- |
| # Samples | Phylotree 17<br>(1 thread) | 1KP3 10k<br>(1 thread) | 1KP3 100k<br>(1 thread) |
| 26 samples | 2 min | 2h 12 min | 4h 33 min |

### Contamination Source

Haplocheck always reports both the major and minor haplotype for each sample. Therefore, possible sources of contamination can be investigated. For example, sample HG00740 from the 1000 Genomes Project shows a contamination level of 2.74% on nDNA (using VerifyBamID2) and 3% on mtDNA (using haplocheck). By looking at the phylogenetic tree that is created for each sample by haplocheck, the contaminating minor haplogroup B2b3a can be identified. The identical haplogroup is also assigned to sample HG01079 that has been analyzed in the same center with a similar mitochondrial copy number. Such phylogenetic information provided within the interactive HTML report can help in identifying the source of contamination for all three types of contamination.

## Discussion

There are many examples in the literature showing the negative impact of artefacts on mtDNA datasets in different areas of research, including medical studies, forensic genetics and human population studies (Bandelt and Salas 2012; He et al. 2010; Ye et al. 2014; Just et al. 2014). The described approach in this paper takes advantage of the mitochondrial phylogeny and is capable of detecting contamination based on mitochondrial haplotype mixtures. By creating several in-silico data sets and analyzing the 1000 Genomes Project data, we show that haplocheck can be used in both targeted mtDNA sequencing studies and WGS studies. We also investigated the influence of the mitochondrial copy number (mtCN) and showed that it must be taken into account when comparing mtDNA to nDNA contamination levels.

Several other methods for contamination detection exist. For nDNA sequences, VerifyBamID2 (Zhang et al. 2020) offers an ancestry-agnostic DNA contamination estimation method and is widely used in WGS studies. Schmutzi (Renaud et al. 2015) provides a contamination estimation tool appropriate for ancient mtDNA. A further approach was suggested in (Dickins et al. 2014), describing a pipeline for contamination detection accessible through the Galaxy online platform (Afgan et al. 2018).

We also identified limitations with the proposed phylogenetic based contamination check in this paper, previously applied in a semi-automatic manner (Avital et al. 2012; Li et al. 2010). There is currently a publication bias in favor of the European mtDNA haplogroups that provides the most phylogenetic details, whereas especially African haplogroups are underrepresented (626 African haplogroups compared to 2,546 European haplogroups in Phylotree 17). While the major changes in the phylogeny were performed during the initial growing process of the tree, the last few years showed only refinements of lineages and branches. Therefore, major changes are no longer expected in the human phylogeny, but data from upcoming sequencing studies will help to refine existing groups. Further, contamination detection based on mitochondrial genomes is limited in scenarios where samples belong to the same maternal line (e.g. mother-offspring). If a contamination between mother and offspring exists, the presented approach is unable to detect it.

Overall, we demonstrated that haplogroup-based analysis as carried out by haplocheck can be used systematically as a quality measure for mtDNA data. Such kind of analysis could become effective prior to data interpretation and publication of mtDNA sequencing projects. Additionally, haplocheck proves to be useful in WGS studies as a fast proxy tool for estimating the nDNA contamination level.

## Software Availability

Haplocheck is available at https://github.com/genepi/haplocheck under the MIT license and requires Java 8 or higher for local execution. All generated data, scripts and reports are available within this repository. The web service can be accessed via https://mitoverse.i-med.ac.at.

## Acknowledgements

We would like to acknowledge the support of the IT Department from the Medical University of Innsbruck (especially Mario Bedenk, Michael Hoertnagl, Matthias Tschugg and Christoph Wild) for providing technical support and resources for the mitoverse web-service.

## Author Contributions

HW and SeS devised the project and implemented the software. LuF developed Cloudgene. SeS, HW, AK and LiF wrote the manuscript. FK and AS supervised the project and contributed to the manuscript. All authors read and approved the final manuscript.

## Footnotes

1 ftp://ftp.1000genomes.ebi.ac.uk/vol1/ftp/phase3/

## References

1000 Genomes Project Consortium, 2015. A global reference for human genetic variation. Nature 526: 68–74.

Afgan E, Baker D, Batut B, van den Beek M, Bouvier D, Cech M, Chilton J, Clements D, Coraor N, Grüning BA, et al. 2018. The Galaxy platform for accessible, reproducible and collaborative biomedical analyses: 2018 update. Nucleic Acids Research 46: W537–W544. http://dx.doi.org/10.1093/nar/gky379.

Albayrak L, Khanipov K, Pimenova M, Golovko G, Rojas M, Pavlidis I, Chumakov S, Aguilar G, Chávez A, Widger WR, et al. 2016. The ability of human nuclear DNA to cause false positive low-abundance heteroplasmy calls varies across the mitochondrial genome. BMC Genomics 17: 1017.

Andrews RM, Kubacka I, Chinnery PF, Lightowlers RN, Turnbull DM, Howell N. 1999. Reanalysis and revision of the Cambridge reference sequence for human mitochondrial DNA. Nat Genet 23: 147.

Avital G, Buchshtav M, Zhidkov I, Tuval Feder J, Dadon S, Rubin E, Glass D, Spector TD, Mishmar D. 2012. Mitochondrial DNA heteroplasmy in diabetes and normal adults: role of acquired and inherited mutational patterns in twins. Hum Mol Genet 21: 4214–4224.

Balciuniene J, Balciunas D. 2019. A Nuclear mtDNA Concatemer (Mega-NUMT) Could Mimic Paternal Inheritance of Mitochondrial Genome. Front Genet 10: 518.

Bandelt H-J, Salas A. 2012. Current Next Generation Sequencing technology may not meet forensic standards. Forensic Science International: Genetics 6: 143–145. http://dx.doi.org/10.1016/j.fsigen.2011.04.004.

Bandelt HJ, Salas A, Lutz-Bonengel S. 2004. Artificial recombination in forensic mtDNA population databases. Int J Legal Med 118: 267–273.

Brandhagen MD, Just RS, Irwin JA. 2020. Validation of NGS for mitochondrial DNA casework at the FBI Laboratory. Forensic Sci Int Genet 44: 102151.

Danecek P, Auton A, Abecasis G, Albers CA, Banks E, DePristo MA, Handsaker RE, Lunter G, Marth GT, Sherry ST, et al. 2011. The variant call format and VCFtools. Bioinformatics 27: 2156–2158.

Das S, Forer L, Schönherr S, Sidore C, Locke AE, Kwong A, Vrieze SI, Chew EY, Levy S, McGue M, et al. 2016. Next-generation genotype imputation service and methods. Nat Genet 48: 1284–1287.

Dayama G, Emery SB, Kidd JM, Mills RE. 2014. The genomic landscape of polymorphic human nuclear mitochondrial insertions. Nucleic Acids Res 42: 12640–12649.

Dickins B, Rebolledo-Jaramillo B, Su MS-W, Paul IM, Blankenberg D, Stoler N, Makova KD, Nekrutenko A. 2014. Controlling for contamination in re-sequencing studies with a reproducible web-based phylogenetic approach. Biotechniques 56: 134–141.

Ding J, Sidore C, Butler TJ, Wing MK, Qian Y, Meirelles O, Busonero F, Tsoi LC, Maschio A, Angius A, et al. 2015. Assessing Mitochondrial DNA Variation and Copy Number in Lymphocytes of ∼2,000 Sardinians Using Tailored Sequencing Analysis Tools. PLoS Genet 11: e1005306.

Fazzini F, Lamina C, Fendt L, Schultheiss UT, Kotsis F, Hicks AA, Meiselbach H, Weissensteiner H, Forer L, Krane V, et al. 2019. Mitochondrial DNA copy number is associated with mortality and infections in a large cohort of patients with chronic kidney disease. Kidney Int 96: 480–488.

Goto H, Dickins B, Afgan E, Paul IM, Taylor J, Makova KD, Nekrutenko A. 2011. Dynamics of mitochondrial heteroplasmy in three families investigated via a repeatable re-sequencing study. Genome Biol 12: R59.

He Y, Wu J, Dressman DC, Iacobuzio-Donahue C, Markowitz SD, Velculescu VE, Diaz LA Jr, Kinzler KW, Vogelstein B, Papadopoulos N. 2010. Heteroplasmic mitochondrial DNA mutations in normal and tumour cells. Nature 464: 610–614.

Huang W, Li L, Myers JR, Marth GT. 2012. ART: a next-generation sequencing read simulator. Bioinformatics 28: 593–594.

Jun G, Flickinger M, Hetrick KN, Romm JM, Doheny KF, Abecasis GR, Boehnke M, Kang HM. 2012. Detecting and estimating contamination of human DNA samples in sequencing and array-based genotype data. Am J Hum Genet 91: 839–848.

Just RS, Irwin JA, Parson W. 2015. Mitochondrial DNA heteroplasmy in the emerging field of massively parallel sequencing. Forensic Sci Int Genet 18: 131–139.

Just RS, Irwin JA, Parson W. 2014. Questioning the prevalence and reliability of human mitochondrial DNA heteroplasmy from massively parallel sequencing data. Proc Natl Acad Sci U S A 111: E4546–7.

Kivisild T, Metspalu M, Bandelt H-J, Richards M, Villems R. 2006. The World mtDNA Phylogeny. Nucleic Acids and Molecular Biology 149–179. http://dx.doi.org/10.1007/3-540-31789-9_7.

Kloss-Brandstätter A, Pacher D, Schönherr S. 2011. HaploGrep: a fast and reliable algorithm for automatic classification of mitochondrial DNA haplogroups. Human. http://onlinelibrary.wiley.com/doi/10.1002/humu.21382/full.

Li M, Schönberg A, Schaefer M, Schroeder R, Nasidze I, Stoneking M. 2010. Detecting heteroplasmy from high-throughput sequencing of complete human mitochondrial DNA genomes. Am J Hum Genet 87: 237–249.

Li M, Schröder R, Ni S, Madea B, Stoneking M. 2015. Extensive tissue-related and allele-related mtDNA heteroplasmy suggests positive selection for somatic mutations. Proc Natl Acad Sci U S A 112: 2491–2496.

Li M, Schroeder R, Ko A, Stoneking M. 2012. Fidelity of capture-enrichment for mtDNA genome sequencing: influence of NUMTs. Nucleic Acids Res 40: e137.

Maude H, Davidson M, Charitakis N, Diaz L, Bowers WHT, Gradovich E, Andrew T, Huntley D. 2019. NUMT Confounding Biases Mitochondrial Heteroplasmy Calls in Favor of the Reference Allele. Front Cell Dev Biol 7: 201.

Renaud G, Slon V, Duggan AT, Kelso J. 2015. Schmutzi: estimation of contamination and endogenous mitochondrial consensus calling for ancient DNA. Genome Biol 16: 224.

Salas A, Schönherr S, Bandelt HJ. 2020. Extraordinary claims require extraordinary evidence in asserted mtDNA biparental inheritance. Forensic Sci. https://www.sciencedirect.com/science/article/pii/S1872497320300454.

Salas A, Yao Y-G, Macaulay V, Vega A, Carracedo A, Bandelt H-J. 2005. A critical reassessment of the role of mitochondria in tumorigenesis. PLoS Med 2: e296.

Samuels DC, Han L, Li J, Quanghu S, Clark TA, Shyr Y, Guo Y. 2013. Finding the lost treasures in exome sequencing data. Trends Genet 29: 593–599.

Schönherr S, Forer L, Weißensteiner H, Kronenberg F, Specht G, Kloss-Brandstätter A. 2012. Cloudgene: a graphical execution platform for MapReduce programs on private and public clouds. BMC Bioinformatics 13: 200.

Van Der Valk T, Vezzi F, Ormestad M, Dalén L, Guschanski K. 2019. Index hopping on the Illumina HiseqX platform and its consequences for ancient DNA studies. Mol Ecol Resour. https://onlinelibrary.wiley.com/doi/abs/10.1111/1755-0998.13009.

Vohr SH, Gordon R, Eizenga JM, Erlich HA, Calloway CD, Green RE. 2017. A phylogenetic approach for haplotype analysis of sequence data from complex mitochondrial mixtures. Forensic Sci Int Genet 30: 93–105.

Weissensteiner H, Forer L, Fuchsberger C, Schöpf B, Kloss-Brandstätter A, Specht G, Kronenberg F, Schönherr S. 2016. mtDNA-Server: next-generation sequencing data analysis of human mitochondrial DNA in the cloud. Nucleic Acids Res 44: W64–9.

Wei W, Pagnamenta AT, Gleadall N, Sanchis-Juan A, Stephens J, Broxholme J, Tuna S, Odhams CA, Genomics England Research Consortium, NIHR BioResource, et al. 2020. Nuclear-mitochondrial DNA segments resemble paternally inherited mitochondrial DNA in humans. Nat Commun 11: 1740.

Wei W, Tuna S, Keogh MJ, Smith KR, Aitman TJ, Beales PL, Bennett DL, Gale DP, Bitner-Glindzicz MAK, Black GC, et al. 2019. Germline selection shapes human mitochondrial DNA diversity. Science 364. http://dx.doi.org/10.1126/science.aau6520.

Yao Y-G, Bandelt H-J, Young NS. 2007. External contamination in single cell mtDNA analysis. PLoS One 2: e681.

Ye K, Lu J, Ma F, Keinan A, Gu Z. 2014. Extensive pathogenicity of mitochondrial heteroplasmy in healthy human individuals. Proc Natl Acad Sci U S A 111: 10654–10659.

Yin C, Liu Y, Guo X, Li D, Fang W, Yang J, Zhou F, Niu W, Jia Y, Yang H, et al. 2019. An Effective Strategy to Eliminate Inherent Cross-Contamination in mtDNA Next-Generation Sequencing of Multiple Samples. J Mol Diagn 21: 593–601.

Yuan Y, Ju YS, Kim Y, Li J, Wang Y, Yoon CJ, Yang Y, Martincorena I, Creighton CJ, Weinstein JN, et al. 2020. Comprehensive molecular characterization of mitochondrial genomes in human cancers. Nat Genet 52: 342–352.

Zhang F, Flickinger M, Taliun SAG, InPSYght Psychiatric Genetics Consortium, Abecasis GR, Scott LJ, McCaroll SA, Pato CN, Boehnke M, Kang HM. 2020. Ancestry-agnostic estimation of DNA sample contamination from sequence reads. Genome Res 30: 185–194.

Zhang R, Wang Y, Ye K, Picard M, Gu Z. 2017. Independent impacts of aging on mitochondrial DNA quantity and quality in humans. BMC Genomics 18: 890.

